# Species Tailoured Contribution of Volumetric Growth and Tissue Convergence to Posterior Body Elongation in Vertebrates

**DOI:** 10.1101/025882

**Authors:** Ben Steventon, Fernando Duarte, Ronan Lagadec, Sylvie Mazan, Jean-François Nicolas, Estelle Hirsinger

**Affiliations:** Department of Developmental and Stem Cell Biology, Institut Pasteur, 25 rue du Doctor Roux, Paris, France; Development and Evolution of Vertebrates, CNRS-UPMC-UMR 7150, Station Biologique, Roscoff, France

**Keywords:** Posterior body elongation, Zebrafish, Mouse, Lamprey, Dogfish, Volumetric Growth

## Abstract

Axial elongation is a widespread mechanism propelling the generation of the metazoan body plan. A widely accepted model is that of posterior growth, where new tissue is continually added from the posterior unsegmented tip of the body axis. A key question is whether or not such a posterior growth zone generates sufficient additional tissue volume to generate elongation of the body axis, and the degree to which this is balanced with tissue convergence and/or growth in already segmented regions of the body axis. We applied a multi-scalar morphometric analysis during posterior axis elongation in zebrafish. Importantly, by labelling of specific regions/tissues and tracking their deformation, we observed that the unsegmented region does not generate additional tissue volume at the caudal tip. Instead, it contributes to axis elongation by extensive tissue deformation at constant volume. We show that volumetric growth occurs in the segmented portion of the axis and can be attributed to an increase in the size and length of the spinal cord and notochord. FGF inhibition blocks tissue convergence within the tailbud and unsegmented region rather than affecting volumetric growth, showing that a conserved molecular mechanism can control convergent morphogenesis, even if by different cell behaviours. Finally, a comparative morphometric analysis in lamprey, dogfish, zebrafish and mouse reveal a differential contribution of volumetric growth that is linked to a switch between external and internal modes of development. We propose that posterior growth is not a conserved mechanism to drive axis elongation in vertebrates. It is instead associated with an overall increase in growth characteristic of internally developing embryos that undergo embryonic development concomitantly with an increase in energy supply from the female parent.

## Introduction

Axial elongation is a widespread mechanism propelling the generation of the metazoan body plan (Martin and Kimelman, 2009). A widely accepted model for posterior body elongation is that of posterior growth, where new tissue is continually added from the posterior tip of the elongating body axis. In vertebrates, a posterior proliferative zone is thought to drive this process by providing new cells to populate the posterior tissues, the mesoderm and spinal cord in particular (Beddington, 1994; Bouldin et al., 2014; Cambray and Wilson, 2002; Mathis and Nicolas, 2000; McGrew et al., 2008; Nicolas et al., 1996; Selleck and Stern, 1991). However, in order to contribute to the elongation of the posterior body, this growth zone must generate a significant increase in volume at the whole embryo level. Therefore, volumetric measurements are required to determine whether this posterior proliferative zone generates sufficient volume increase to elongate the posterior body. Several lines of evidence suggest that cell proliferation is not strictly required for axis elongation. The zebrafish *emi1* mutant, in which cell divisions are blocked from the beginning of gastrulation (Riley et al., 2010; Zhang et al., 2008) is not truncated but only shortened by around 30%. Blocking of cell division with the use of mitomycin C and aphidicolin failed to affect elongation rate in the chick embryo (Bénazéraf et al., 2010). Furthermore, a role for proliferation in driving elongation has been challenged in other morphogenetic processes, for example during mouse limb bud outgrowth (Boehm et al., 2010).

In addition to cell proliferation, several other cell behaviours may contribute to growth, (i.e. an increase in tissue volume). A model has been proposed in the chick whereby a gradient of random cell motility generates elongation by decreasing cellular density within the caudal region of the presomitic mesoderm (PSM) (Bénazéraf et al., 2010). In addition, several other cell behaviours such as cell swelling or an increase in extra-cellular matrix production may also be contributing to growth. To incorporate all potential cellular behaviours that could contribute to growth, morphometric measurements at the whole structure level are required to assess degree of volume increase that may or may not be concurrent with axis elongation.

In addition to volumetric growth, several non-growth cell behaviours have also been proposed to drive elongation via tissue convergence, particularly in anamniote embryos such as zebrafish. Gastrulation-like cell rearrangements such as convergence-extension, together with novel cellular movements have been observed in zebrafish posterior elongation (Kanki and Ho, 1997). Also, blocking FGF signalling leads to an inhibition of cellular flow in the tailbud and the disruption of axis elongation (Lawton et al., 2013). In addition, as the presumptive territory for the posterior body has not yet been mapped, it could be that further convergence of cells from domains lateral to the tailbud also contributes to the elongation of the posterior axis. However, while these studies point for a clear role of cell rearrangements, they do not rule out an additional role for volumetric growth during axial elongation in zebrafish.

We aim to determine the relative contribution of volumetric growth vs. tissue deformation (i.e. elongation in the absence of volume increase) to posterior body elongation. As the zebrafish embryo develops externally, with only a limited energy supply in the form of a relatively small yolk sack, a second aim is to assess the degree of growth that is concomitant with posterior body elongation internally developing mouse and dogfish embryos that develop with increased maternal energy supply compared with externally developing zebrafish and lamprey embryos.

We first generated a fate map of the zebrafish posterior body and found a considerable contribution of cells directly into both the spinal cord and PSM without having first transitioned through the tailbud. Our morphometric analysis conducted throughout elongation at different length-scales (i.e. whole posterior body, segmented vs unsegmented regions and individual tissues) revealed that axis elongation occurs via tissue convergence, initially in the absence of volume increase. The photolabelling and tracking of the unsegmented region shows that there is no posterior growth in zebrafish elongation. Whole structure and tissue specific photolabels then revealed that growth does occur at later stages within the notochord and spinal cord. FGF inhibition blocks tissue convergence within the tailbud and PSM rather than controlling posterior growth. Comparing two internally developing (mouse and dogfish) and two externally developing vertebrates (the zebrafish and lamprey), we find that posterior growth is not a conserved process in vertebrates, but is instead correlated with an internal mode of development.

## Results

### The prospective posterior body is not restricted to the tailbud region

In the zebrafish, the embryonic axis is already established by the late gastrula with the head at the anterior pole. At this stage the prospective anterior trunk somites (1-12) are already present within the unsegmented pre-somitic mesoderm (Kanki and Ho, 1997; Kimmel et al., 1995). Subsequently, the remaining 20 segments of the body form in an anterior to posterior fashion through the process of posterior axis elongation. Previous studies analysing the role of cell movements in driving axial elongation in the zebrafish have focused on cells within the tailbud (Kanki and Ho, 1997; Lawton et al., 2013). However the limits of the posterior body territory are unknown from the end of gastrulation up to the 12-somite stage. We therefore mapped the limits of the prospective posterior body between the end of gastrulation and 12-somite stage. We generated 12 independent movies in which clusters of 10-20 cells where photolabeled with the use of a nuclear localised KiKGR construct (S1 Movie), a photoconvertible fluorescent protein that switches from green to red upon UV exposure (Hatta et al., 2006). This allowed us to track these cells and determine their fate with respect to the posterior body and tailbud. We observed that cells coming from domains lateral to the tailbud converge and are incorporated into the tailbud (blue; Fig. 1A-C). In addition, we found that many cells entered directly into both the spinal cord (yellow; Fig 1A-C) and pre-somitic mesoderm (PSM; red; Fig. 1A-C) without having passed through the tailbud region (blue; Fig. 1A-C) (S1 Movie). This suggests that the prospective posterior body region at late gastrula stages is much larger than previously thought and not restricted to the tailbud region, which prompted us to examine the degree to which growth (i.e. volume increase) versus convergence is contributing to the elongation of the posterior body axis in zebrafish.

**Figure 1.**
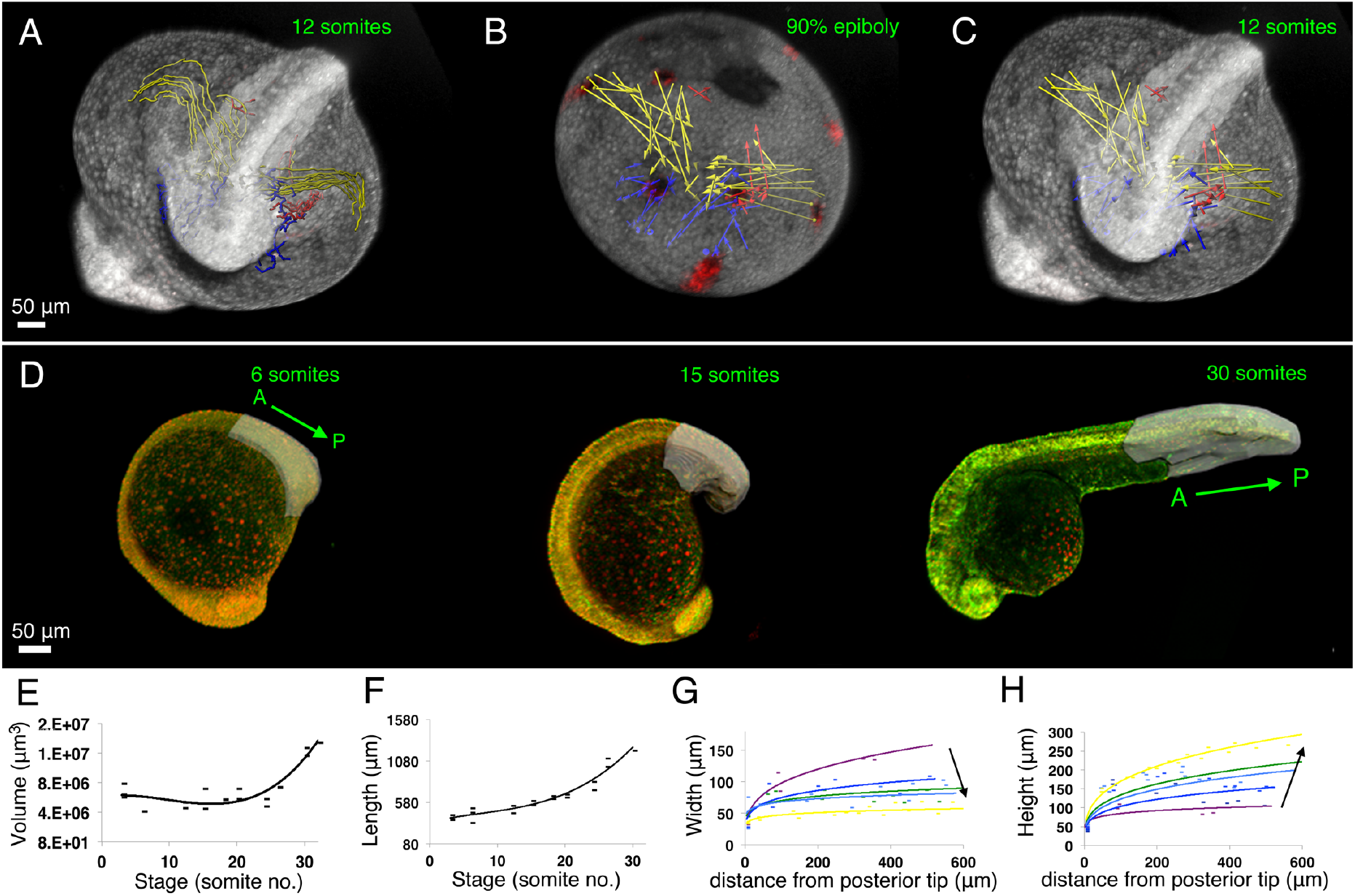
Posterior axis elongation in zebrafish occurs firstly in the absence of growth followed by a later growth phase. (A-C) Tracks (A) and displacement vectors (B,C) of spinal cord (yellow) tailbud (blue) and PSM (red) precursors from automated tracking of photolabeled nuclear from the 90%epiboly to the 12 somite stage in order to determine lateral edges of the prospective posterior body axis. (D) Taking the 12^th^ somite as the trunk/tail boundary, surface reconstructions (grey region) were created at consecutive stages through axial elongation. Embryos shown in lateral view with A and P marking anterior and posterior poles of the posterior body. (E-H) Plots of morphometric measurements of volume (E; polynomial fit, r^2^= 0.83, n=21) and length (F; polynomial fit, r^2^= 0.94, n=21) against time (expressed as total number of somites formed). (G-H) Plots of width (G) and height (H) against distance from posterior tip of the tail to where each measurement was taken (colour code: purple= 12 somites (s) (n=5), dark blue= 15s, light blue= 18s (n=7), green= 20s (n=12), yellow= 24s (=17). Data points show individual measurements, n=total number of measurements from 3 embryos per stage.

### Posterior axis elongation in zebrafish occurs through extensive convergence, initially in the absence of growth followed by a later growth phase

To decipher the modalities of zebrafish posterior body elongation, we performed a dynamic morphometric analysis at the whole structure level. Using this early fate map and samples fixed from 12-somite stage on, we built 3D surface reconstructions of the posterior body throughout the process of axis elongation (Fig. 1A; S2 Movie). From these, we extracted quantitative information on volume, length, width and height changes. Surprisingly, despite a continuous elongation of the posterior body from early stages (Fig 1C), we did not observe an increase in volume until the 24-somite stage (Fig 1B). To generate axis elongation in the absence of growth, tissues must decrease in either width or height at early stages. We observe that during the initial stages of posterior body elongation (purple to yellow lines in Fig. 1D, E), there is a four-fold decrease in width (Fig. 1D). This contrasts with the posterior body height, which increases throughout elongation, suggesting that convergence is contributing both to an increase in length and height (Fig 1E). These results demonstrate that whilst growth may contribute to axial elongation from the 24 somite stage onwards, much of the early stages of posterior body elongation are driven by convergence.

### The zebrafish posterior axis does not elongate by posterior growth

We next wanted to determine in zebrafish whether the posterior axis elongates by addition of tissue at its posterior end (i.e. posterior growth). Such a mode of elongation implies that the volume of tissue generated by the posterior tip should increase over time. We thus labelled the posterior tip of the embryonic axis that has not yet undergone segmentation by somitogenesis (the unsegmented region) and followed its growth over time. We took embryos previously injected with mRNA encoding cytoplasmic Kikume Green to Red (KikGR). Illuminating the unsegmented region with UV at the 15-somite stage results in the labelling of all cells within this region and their descendants. We then performed time-lapse imaging of the unsegmented region photolabels (Fig. 2A, S3 Movie). We tracked the photolabels for a period of five hours, as at later stages unsegmented derivatives have progressed significantly into the segmented portion of the body axis. Automatic thresholding of the photoconverted signal allows us to obtain continuous morphometric information on the relative volume and shape changes of these regions and their derivatives over time. As cells are exiting the tailbud and entering the PSM, we saw considerable mixing of labelled cells with unlabelled cells of the rostral PSM. As this cell mixing may contribute to elongation via cell intercalation, we took for our length measurements the most anterior photolabeled cell. For volume measurements, we segmented and measured the photoconverted (i.e. red) signal only. Although the unsegmented region of the posterior body contributes to elongation (unsegmented: 2.67 fold increase; Fig. 2B), this region does not increase in volume (0.98 fold change; Fig. 2C). In order to determine whether the unsegmented region contributes any growth to posterior body elongation during the growth phase, we photolabelled at the 21-somite stage and measured its morphogenesis through to the completion of somitogenesis (Fig. 2D). This confirmed the absence of unsegmented region growth during the late stages as well (Fig. 2F). However, this region continues to contribute to posterior body elongation at these late stages, although to a lesser extent (Fig. 2E; 1.61 fold increase). Thus, no tissue growth from the unsegmented region is observed during posterior body elongation in zebrafish.

**Figure 2.**
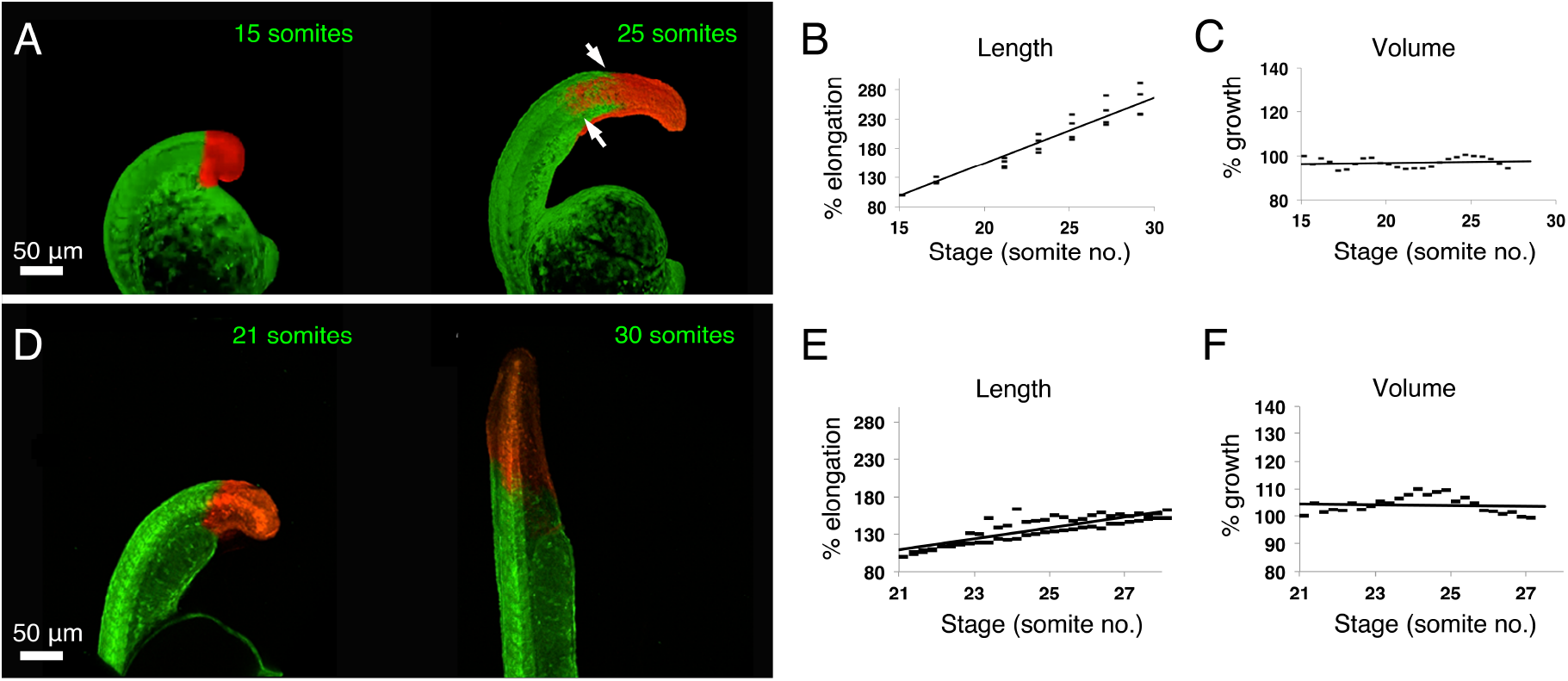
The zebrafish posterior axis does not elongate by posterior growth. (A, D) Stills from time-lapse movies of embryos injected at the one cell stage with Kikume mRNA and regions of the posterior body photo-labelled in the unsegmented at the 15 somite (A) or the 21 somite stage (D). All embryos shown in lateral view with posterior to the top and anterior to the bottom. (B, E) Photoactivated region length (% of final length) plotted against time (number of somites formed), n(B)=25, n(E)=51. (C, F) Photoactivated region volume (% of final volume) plotted against time (number of somites formed), n(C)=25, n(F)=25. Note that photolabels in the unsegmented region do not increase in volume at either stage (C,F). White arrows in (A) indicate the displacement of labels within the somitic mesoderm (lower arrows) relative to the spinal cord (upper arrows). Data points show individual measurements, n=total number of measurements from 3 embryos per experiment.

### The segmented region contributes to elongation via both lengthening and growth

During posterior body elongation, growth is happening at the whole structure level but not in the unsegmented region. We thus tested whether the segmented region exhibited growth. We performed again the photolabel experiment described in Fig. 2 but this time at the level of the segmented region. Unlike the unsegmented region, the segmented region does undergo both elongation (2.19 fold increase, Fig. 3B) and growth (1.25 fold increase, Fig. 3C). Anterior growth occurs in already segmented regions where tissue differentiation is occurring. Importantly, these differences in growth between unsegmented and segmented regions are mirrored by a rapid decrease in cell proliferation within the unsegmented region relative to more anterior structures (Bouldin et al., 2014); (S4 Figure). Overall, these indicate that the segmented region of the body axis grows at least in part by increasing its cell numbers whereas the unsegmented mesod’erm elongates via cell rearrangements in the absence of tissue growth.

**Figure 3.**
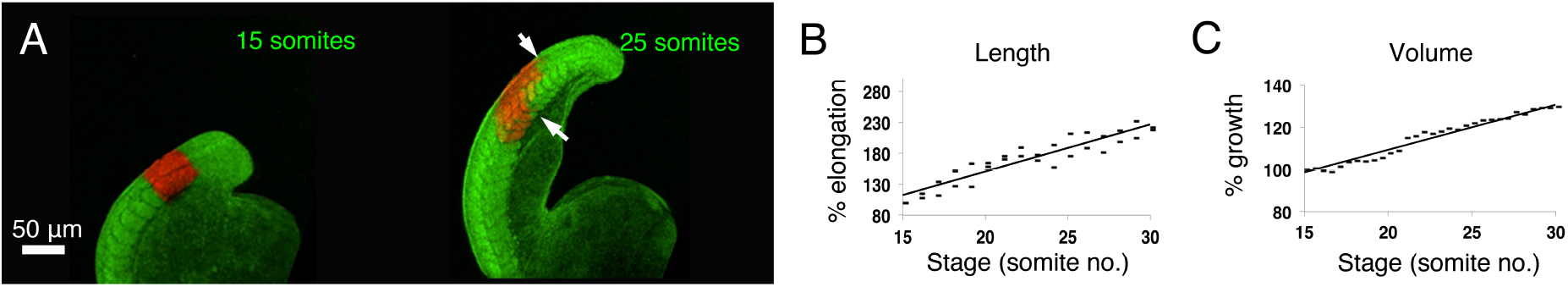
The segmented region contributes to elongation via both lengthening and growth. (A) Stills from time-lapse movies of embryos injected at the one cell stage with KikGR mRNA and the segmented region of the posterior body photo-labelled at the 15 somite somite stage. All embryos shown in lateral view with posterior to the top and anterior to the bottom. (B) Photoactivated region length (% of final length) plotted against time (number of somites formed; n(B)= 29). Photoactivated region volume (% of final volume) plotted against time (number of somites formed; n(C)=31. White arrows in (A) indicate the displacement of labels within the somitic mesoderm (lower arrows) relative to the spinal cord (upper arrows). Data points show individual measurements, n=total number of measurements from 3 embryos per experiment.

### Differential tissue contributions to both growth and convergence and extension

We noticed that regions photolabelled in the unsegmented domain progressed further anteriorly into the pre-somitic mesoderm (Fig. 2A; lower white arrow) as compared to into the spinal cord (Fig. 2A; upper white arrow). The converse was observed for labels in the segmented region, which result in spinal cord cells extending more posteriorly into the tail than cells that were labelled in the somites (Fig. 3A; white arrows). These observations are suggestive of differential contributions of both spinal cord and somitic tissues to growth and/or convergence and extension. To test this further, we photo-labelled small regions of tissue within the tailbud, the pre-somitic mesoderm (PSM), somites and spinal cord (Fig. 4A-C). This analysis allows us to make a quantitative comparison of the contribution of these tissues to growth by plotting the percentage volume increase over time (Fig. 4D). In addition, we analysed the contribution of each tissue to convergence and extension by plotting the length:width ratio against time for each structure (Fig. 4E). The spinal cord contributes the most to growth (Fig. 4B, D, blue). Importantly, cell proliferation is maintained in the spinal cord throughout elongation (S4 Figure). This tissue also has a large positive increase in the length:width ratio (Fig 4B,E; S5 movie; blue), suggesting that either growth is anisotropic (through oriented cell division or oriented growth of the cells) or that additional cell rearrangements such as convergence-extension are leading to a thinning of this structure. The cells in transit from the tailbud to the PSM contribute the most to thinning and lengthening. They undergo the most dramatic increase in length:width ratio, but in the absence of growth (Fig 4C-E; S6 movie; green). Once cells have entered the PSM however, little further thinning and lengthening occurs (Fig. 4B,E; S5 movie; yellow). This tissue also undergoes a slight compaction during somite formation (Fig. 4B,D; yellow).

**Figure 4.**
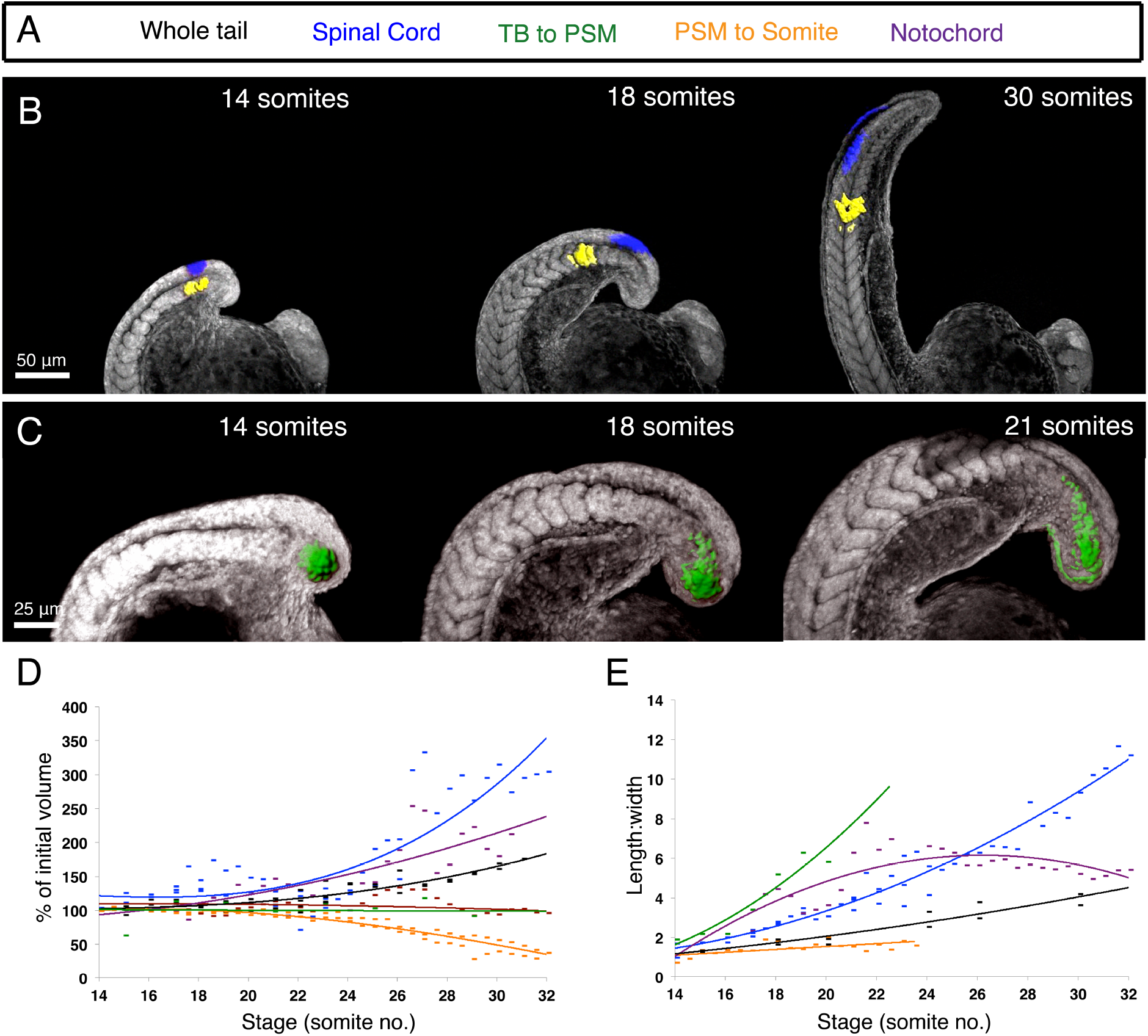
The spinal cord and notochord are the principal contributors to anterior growth and the TB to PSM transiting cells to thinning and lengthening. (A) Different tissues are colour coded according to the key shown. (B) Stills from time-lapse movies of embryos injected at the one cell stage with a Kikume and photo-labelled in the spinal cord and as cells transition from the PSM to newly formed somites. (C) Cells transitioning between the tailbud (TB) and pre-somitic mesoderm. (D, E) Plots of volume (D; n(Spinal cord)=51, n(notochord)=31, n(PSM to somites)=58, n(TB to PSM)=23) and length:width (E; n(Spinal cord)=43, n(notochord)=30, n(PSM to somites)=17, n(TB to PSM)=12) of each photo-labelled region against time (expressed as total number of somites formed). Data points show individual measurements, n=total number of measurements from 3 embryos per experiment.

In addition to growth in the spinal cord, it is known that inflation of the notochord by the formation of fluid-filled organelles has an important role in the elongation of the zebrafish axis (Ellis et al., 2013). In line with this, we observe a considerable degree of volume increase in notochord photolabels (Fig. 4D). This is mirrored by the increase in the volume of bounding-boxes surrounding notochord photo-labels (Fig. 4D; purple; includes the organelle volume) without a corresponding increase in the volume of the KikGR-labelled cytoplasm (Fig. 4D; magenta; excludes the organelle volume). In addition to inflation and proliferation, the notochord also undergoes convergence and extension (Glickman et al., 2003), which is mirrored by an increase in length:width (Fig. 4E; purple). However, as cells begin to inflate, they do so in all directions, leading to a later decrease in length:width ratio (Fig. 4E; purple).

The impact of these distinct tissue deformations on the elongation of the body axis as a whole will depend on their initial size with respect to the whole body axis, as a large volume increase in a small tissue may not contribute much to the morphogenesis of the whole structure. To investigate this, we segmented the paraxial mesoderm, spinal cord and notochord at the 15-somite stage and measured their volume as a proportion of the posterior body volume (S7 Figure). The paraxial mesoderm makes up the largest portion of the axis, at approx. 23% (S7 Figure), thus the convergence of this tissue is likely a major contributor to axis elongation. The spinal cord also forms a significant proportion of the zebrafish axis (12%; S7 Figure) and therefore growth and convergence of this tissue is an additional major contributor to axis elongation. The notochord, at 1.9% of the whole posterior body axis is also likely to contribute although to a much lesser extent than the other tissues. The remaining approx. 63% of the posterior body consists of inter-tissue components, the non-neural ectoderm and endodermal tissues which all may be contributing to differing extents to axis elongation.

### FGF signalling is only required for convergence and extension, not growth of the zebrafish posterior body

FGF has been shown to be important for zebrafish posterior elongation [14] and studies in amniotes suggest that FGF is required to generate posterior growth (Bénazéraf et al., 2010; Bouldin et al., 2014; del Corral and Storey, 2004; Olivera-Martinez et al., 2012). Therefore, we applied our morphometric method to test the role for FGF in generating elongation in zebrafish where there is no posterior growth. To circumvent effects on mesoderm induction and patterning as a consequence of inhibiting FGF signalling prior to the end of gastrulation, we made use of known inhibitors of FGF receptors: SU5402 and PD173074. Addition of these drugs at the 10-somite stage and examination at the 32-somite stage lead to clear affects on posterior axis morphology (Fig. 5A-D), however segments were still present, allowing us to perform surface 3D surface renderings of the posterior axis region. Neither treatment affected posterior axis volume when compared to DMSO treated controls (Fig. 5E, p>0.7 in all cases). However, we did see a significant reduction in axis length (Fig. 5F, p<0.01). This coincided with an increase in mean width (Fig. 5G, p<0.01), suggesting that an inhibition of convergence and extension is the principle cause of axis shortening upon inhibition of FGF signalling. To test this we next repeated our small photo-labellings of the tailbud in both embryos cultures in DMSO (Fig. 5H) and together with PD173074 (Fig 5I) and monitored their transition into the unsegmented region by time-lapse microscopy. No effect was observed on the volume of the tailbud photo-labelled cells upon addition of PD173074 (Fig. 5J), however we did observe an inhibition of the length:width ratio increase that is normally observed for these cells (compare red to green lines in Fig. 5K). Taken together, these results demonstrate that the principle role of FGF in posterior axis elongation in the zebrafish is to control tissue convergence as cells exit the tailbud and enter the PSM.

**Figure 5.**
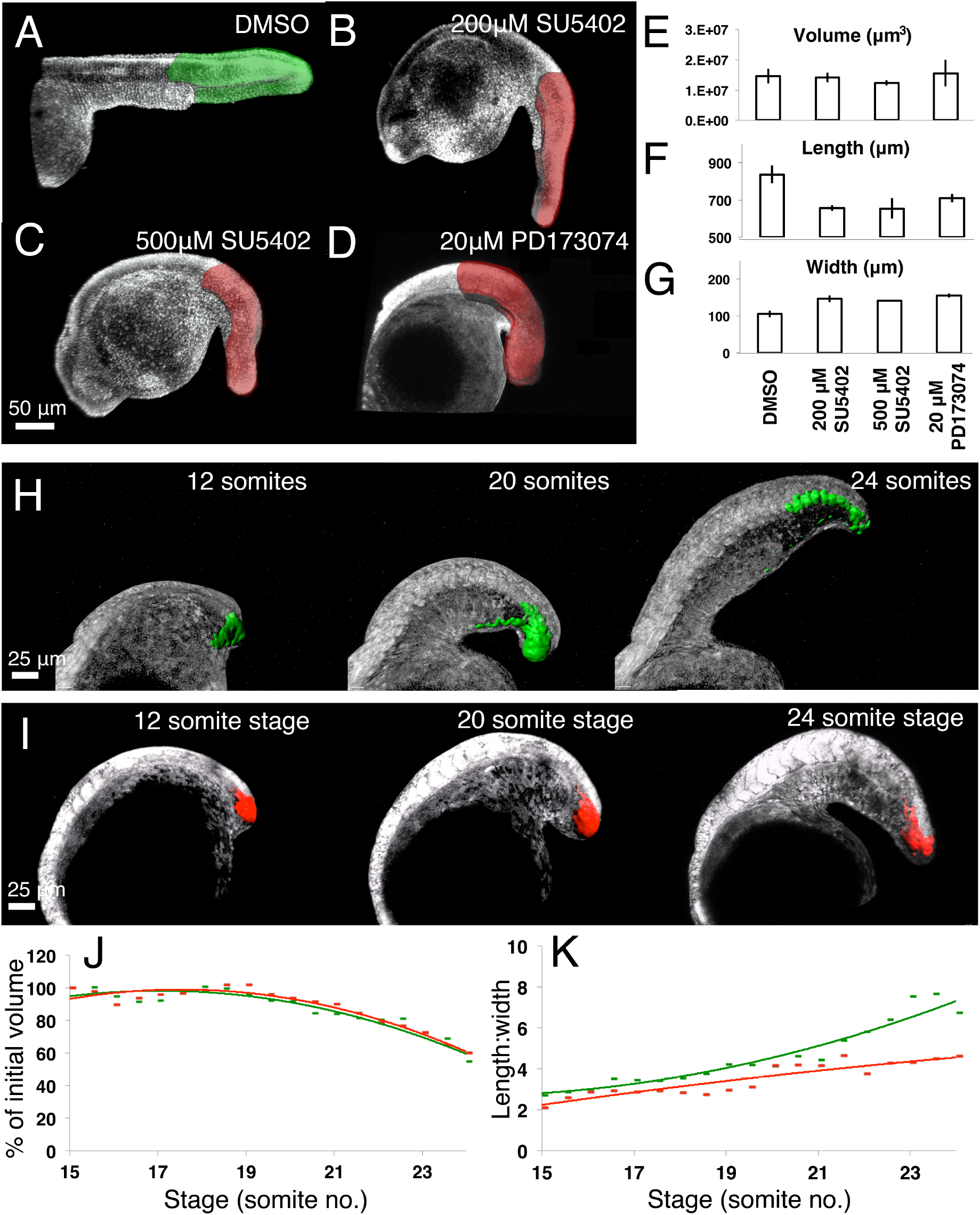
FGF signalling is required for thinning and lengthening not growth of the zebrafish posterior body. (A-D) Embryos were incubated in either 500μM DMSO (A), 200μM SU5402 (B), 500μM SU5402 (C) or 20μM PD173074 (D) from the 10 somite stage until the 32 somite stage and imaged by confocal microscopy. This allowed for the segmentation of the posterior axis and measurement of volume (E), length (F) and width (G) for each condition. (H-I) Stills from time-lapse movies of embryos in which smalls regions of cells within the tailbud are photolabelled and tracked until their entry into the somitic mesoderm in the presence of 500μM DMSO (H) and 20μM PD173074 (I). This allowed for the quantification of volume (J) and length:width (K) over time. Green lines correspond the control situation, red lines indicate treatment with PD173074.

### Posterior growth is not a conserved developmental mechanism across vertebrates

The absence of posterior growth in zebrafish is striking and prompted us to examine the degree of volume increase that occurs in unsegmented vs. segmented portions of the body axis in a wider range of vertebrates (Fig. 6E). In particular, we were interested to determine whether the lack of posterior growth in zebrafish was characteristic of the external development of anamniote embryos. Therefore we compared our morphometric measurements in zebrafish (Fig. 6C) to the amniote mouse embryo (Fig. 6D) and to a basal anamniote vertebrate, the lamprey (Fig. 6A). As a further comparison, we analysed the anamniote dogfish that is evolutionarily basal to teleost fish (Fig. 6E). Importantly however, these embryos develop internally within an egg case and together with a large yolk supply. Thus, posterior growth in this animal would argue for a direct relationship between an increased maternal energy supply and posterior growth, rather than for a later evolution of posterior growth in amniotes.

**Figure 6.**
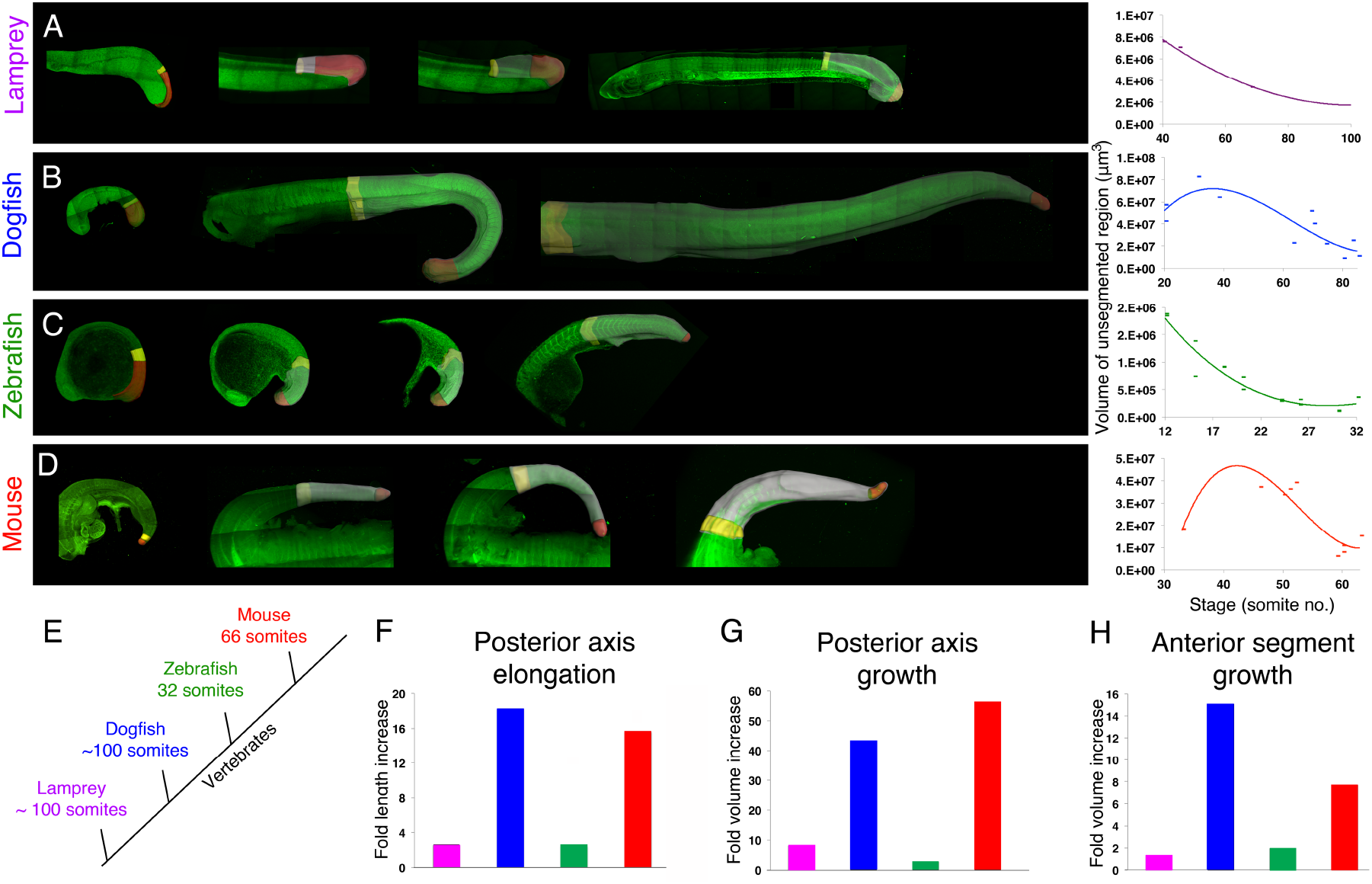
Comparative 4D morphometric analysis of posterior body elongation across vertebrates. (A-D) Maximal projections of tiled z-stacks of lamprey (A, n=7), dogfish (B, n=11), zebrafish (C, n=15) and mouse (D, n=10) embryos at consecutive stages through posterior body elongation. All embryos shown in lateral view with posterior to the right. Grey regions show the entire segmented posterior body, yellow regions show the two most anterior body segments and the red region shows the posterior-most unsegmented region. Corresponding plots of unsegmented region volume over time (in somite number) are shown to the right. (E) Simplified phylogeny for each species studied together with total somite number. (F) Fold change in length (bars, left hand scale) and elongation rate (fold length increase/time (hours); crosses, right hand scale) for each species. (G,H) Fold change in volume of whole posterior body axis and anterior segments (H). Purple bar= lamprey, blue= dogfish, green= zebrafish, red= mouse.

Both dogfish and mouse embryos showed an initial increase in unsegmented region volume at early stages that is consistent with the production of additional tissue from a posterior growth zone (Fig. 6B,D). As expected for the lack of unsegmented region growth in zebrafish, this region undergoes a continual reduction in volume throughout the elongation of the posterior axis, as tissue is continuously being segmented in the anterior with no additional tissue being produced from the tailbud (Fig. 6C). A similar continual reduction is observed in lamprey, suggestive of a null or limited contribution of unsegmented region growth in this basal vertebrate (Fig. 6A). The unsegmented region growth in mouse and dogfish is mirrored by a high level of axis elongation (Fig. 6F) and overall growth (Fig. 6G).

To compare the degree to which segmented region growth contributes to axial elongation, we measured the volume of the two most anterior segments between species (shaded yellow in Fig. 6A-D). This revealed that anterior growth is occurring in all species, albeit at different intensities that correlate with the intensity of the overall growth and elongation (Fig. 6H). Importantly, the fold increase in anterior segments is much lower than that of the overall posterior body (15.0 compared to 40.6 for dogfish; 7.7 compared to 56.1 for mouse), suggesting that much of the overall growth in these two organisms is indeed produced from the posterior unsegmented tissue. Taken together, these results demonstrate that whilst unsegmented region growth plays a major role in the elongation of the posterior body axis of both mouse and dogfish embryos, posterior body elongation in both lamprey and zebrafish embryos occurs largely in the absence of unsegmented region growth.

## Discussion

### Multi-tissue contribution to posterior body elongation in zebrafish

Our multi-scalar morphometric analysis allows for several conclusions to be drawn relating to the differential tissue contribution to posterior axis elongation in zebrafish. Firstly, whilst no additional volume is generated from the posterior unsegmented portion of the axis, convergence of this region is observed as cells transit from the tailbud into the PSM. Secondly, volume increase in the anterior segmented portion of the axis is correlated with an increase in the size of both the spinal cord and notochord. The observation that the anterior segmented region of the axis may be a major contributor to axial elongation explains the bi-phasic growth curve that is observed at the level of the structure as a whole (Fig. 1B). At early stages, the spinal cord and notochord growth, although occurring in anterior trunk structures, have not yet begun to occur within the posterior body. However, convergence and extension of cells entering the PSM is well underway, resulting in a thinning and lengthening of the posterior body axis in the absence of growth (Fig. 1B,E). At later stages, the dual processes of spinal cord growth and notocord inflation have reached the posterior body, resulting in an overall increase in posterior body volume (Fig. 1B,E) and a relative displacement of somitic and spinal cord cells (Fig. 2A; white arrows). Additional cell behaviours such as cell rearrangement, cell shape change and orientated cell division must act together with this growth in order to drive the elongation of the embryonic axis. While our morphometric measurements preclude conclusions to be drawn at the cellular level, they do provide a framework in which to incorporate such observations and to enable the construction of a complete model of this complex morphogenetic process. Once this is attained, it may then be possible to inhibit distinct cellular behaviours and to determine their role in driving axial elongation.

### Origin of cells that make up the posterior body

We demonstrate that posterior growth is not occurring during zebrafish posterior body elongation. The concept of posterior growth is often linked to the presence of a tailbud-resident and self-renewing progenitor population (Beddington, 1994; Cambray and Wilson, 2002; Mathis and Nicolas, 2000; McGrew et al., 2008; Nicolas et al., 1996; Selleck and Stern, 1991) that generate the cells that will make up the posterior body. These cells have been proposed to exist in the zebrafish tailbud (Martin and Kimelman, 2012). Considering the lack of expansion of the unsegmented region, our results are consistent either with the absence of such cells or that they have limited impact on the process of posterior body elongation. In support of this is the *emi* mutant phenotype that shows an axis shortening and not a truncation (Riley et al., 2010; Zhang et al., 2008). In addition, the cell transplantation experiments reported in (Martin and Kimelman, 2012) did not generate long tailbud-anchored clones as would be expected from tailbud-resident, self-renewing stem cells (Tzouanacou et al., 2009). The existence of tailbud-resident, self-renewing stem cells thus awaits long-term clonal analysis, similar to what has been performed in the mouse (Tzouanacou et al., 2009). Consistent with the absence of significant growth contribution during axis elongation, our fate mapping shows that a large proportion of the cells that will make up the posterior body come from regions lateral to the tailbud, and not from proliferation of tailbud cells.

### Posterior growth is associated with internal developmental mode

The null or limited posterior growth in zebrafish and lamprey is in stark contrast to both the mouse and dogfish where we observe large amounts of volume increase in both unsegmented and segmented regions of the body axis. The dogfish embryo develops inside an egg case with a large yolk supply that is reminiscent of avian embryos (Sauka-Spengler et al., 2003), and the mouse embryo develops together with a large energy supply from the placenta. This is in contrast to the larval-feeding zebrafish that must first establish a full complement of posterior somites in order to swim and find food to grow. The prolarval lamprey also swims, before later undergoing a transition to filter feeding from the sediment in which they are buried during an extensive larval period. Therefore, growth appears to be associated with the increase in energy (either placental in the case of mammals or in the form of a large yolk supply in the case of birds and dogfish) that arises from an internal developmental mode. Whilst shape change behaviours are likely to contribute in all cases, an internal mode of development may have allowed for increased contribution of growth behaviours in embryos such as mouse and dogfish.

Despite these differences, there is a conservation of the role of both FGF and Wnt signaling pathway components and downstream Cdx and Brachyury transcription factors for posterior axis elongation across a range of vertebrates (Baker et al., 2010; Esterberg and Fritz, 2009; Marlow et al., 2004; Row and Kimelman, 2009; Shimizu et al., 2005; Thorpe, 2005; Yang and Thorpe, 2011). *Brachyury* is also expressed in the tailbud during axis elongation in dogfish and lamprey embryos, which is further suggestive of a conservation in the expression of core posterior growth zone markers across vertebrates (Sauka-Spengler et al., 2003). Furthermore, *Brachyury, Wnt3*, *Wnt5* and *Wnt6* are present within the tailbud of amphioxus embryos (Schubert et al., 2001), considered to be the closest living relative of the chordate ancestor (Schubert et al., 2006). A growth zone driving body elongation has also been identified in several invertebrate embryos where it has been shown to be associated with Wnt signaling in spiders (McGregor et al., 2008), Artemia (Copf et al., 2004) and Tribolium (Bolognesi et al., 2008; Copf et al., 2004; Schulz et al., 1998). Altogether these observations have led to the proposal of the existence of a universally conserved posterior growth zone across metazoans (Martin and Kimelman, 2009). However, our results suggest that posterior growth is not a conserved mechanism to drive axis elongation in vertebrates. Thus, it is likely that observed conserved molecular mechanisms control different cellular behaviors according to different modes of elongation: they are required to direct cell fate decisions and/or to regulate cellular flow as cells exit the tailbud and/or to maintain a posterior growth zone. In the case of zebrafish, FGF signalling has switched from a role in posterior growth zone maintenance to a role in controlling the cell movements. This is supported by the observations that FGF is required to maintain cellular flow in zebrafish (Lawton et al., 2013) and that a late activation of a dominant negative FGF receptor does not result in the loss of *ntl* expression (Martin and Kimelman, 2010).

### Supplementary data

**S1 Movie. Example movie to determine posterior and lateral limits of the prospective posterior body.**

**S2 Movie. Example 3D reconstructions of the posterior body.**

**S3 Movie.**
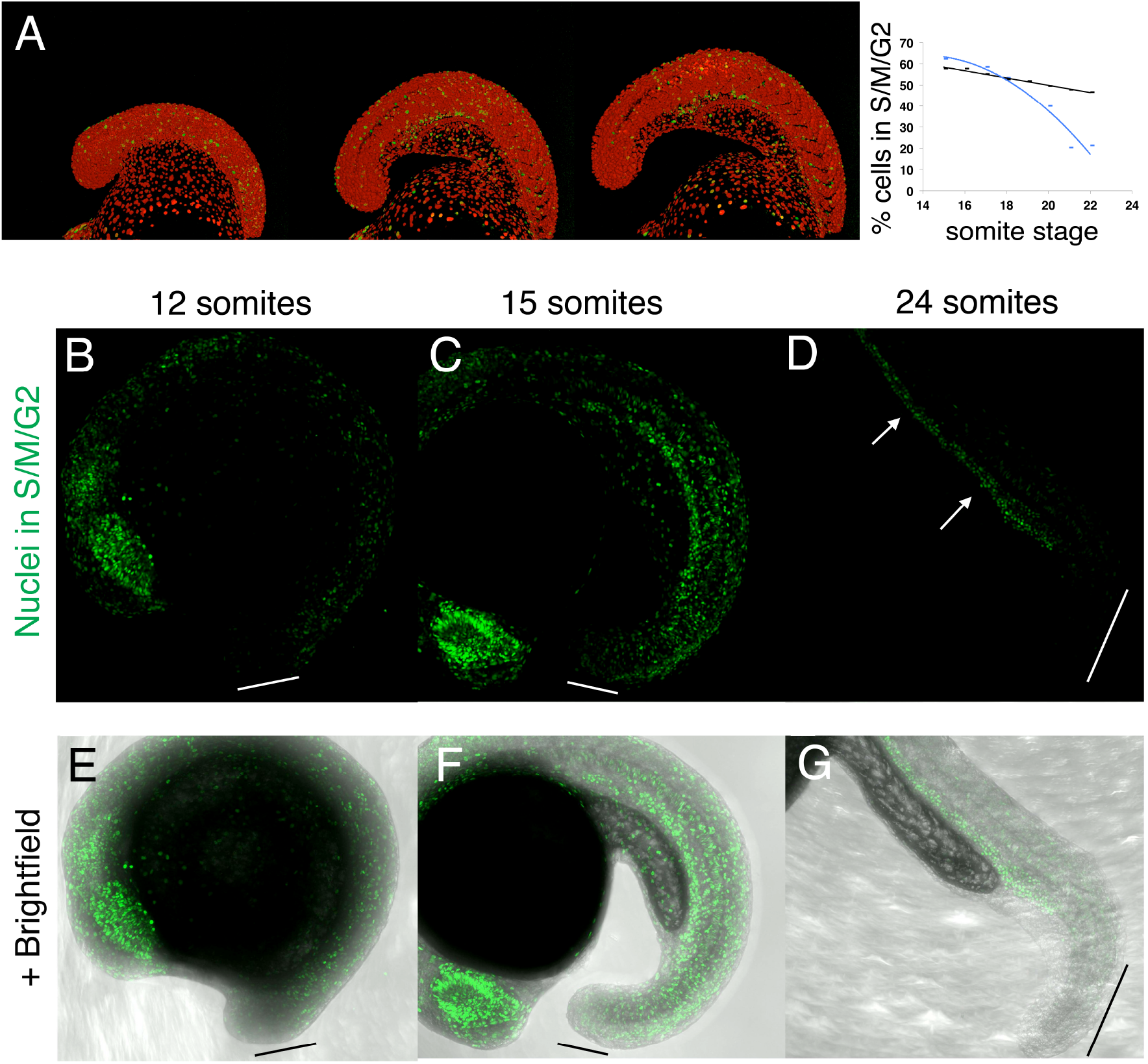
Example of photolabelling in the unsegmented region of the posterior body

**S4 Figure. Cell divisions decrease rapidly in the tailbud region but continue into late stages in anterior structures.** A) Stills from a move of Fucci embryos (Sugiyama et al., 2009) injected with nuclear mCherry to label all nuclei. Quantification of green nuclei vs. total nuclei as marked by mCherry reveals a gradual decline in cell divisions within the posterior body as a whole (black line), however cell divisions in the tailbud decrease much more rapidly (blue line). (B-G) Maximal projections of Fucci green embryos fixed at the 12 (B,E), 15 (C,F) and 24 (D,G) somite stages and imaged by confocal microscopy. Very few cells in S/M/G2 can be observed in the tailbud throughout the process of posterior body elongation (white lines indicate tailbud position in (B-C)). However cells continue to divide in more anterior structures. By the 24 somite stage, cell divisions are mostly restricted to blood precursors within the dorsal aorta (D, white arrows).

**S5 Movie. Example of photolabelling in spinal cord and pre-somitic mesoderm**

**S6 Movie.**
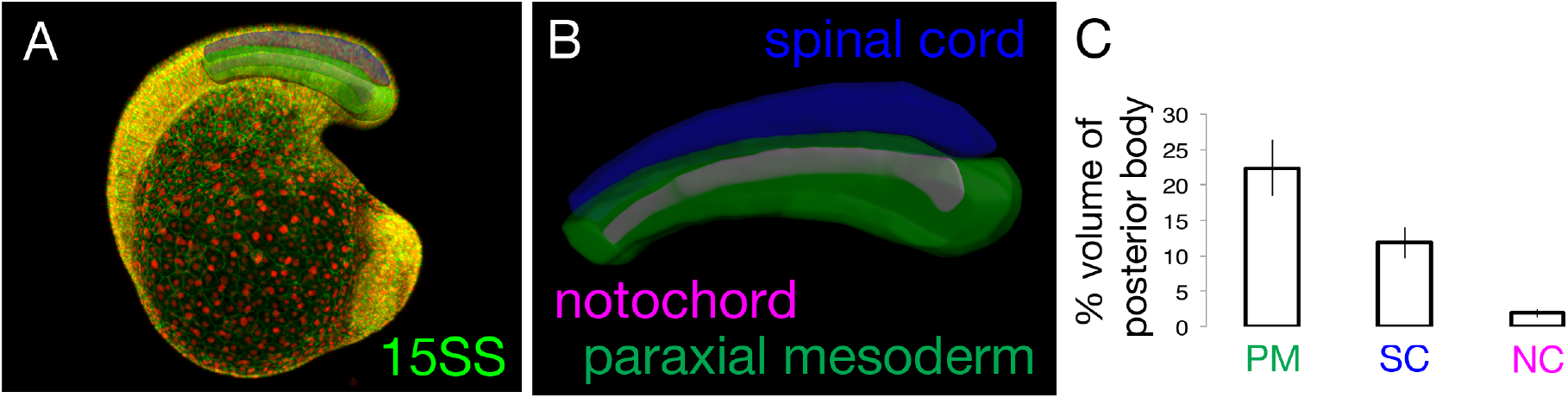
Example of photolabelling in tailbud as cells transition to the pre-somitic mesoderm

**S7 Figure.**
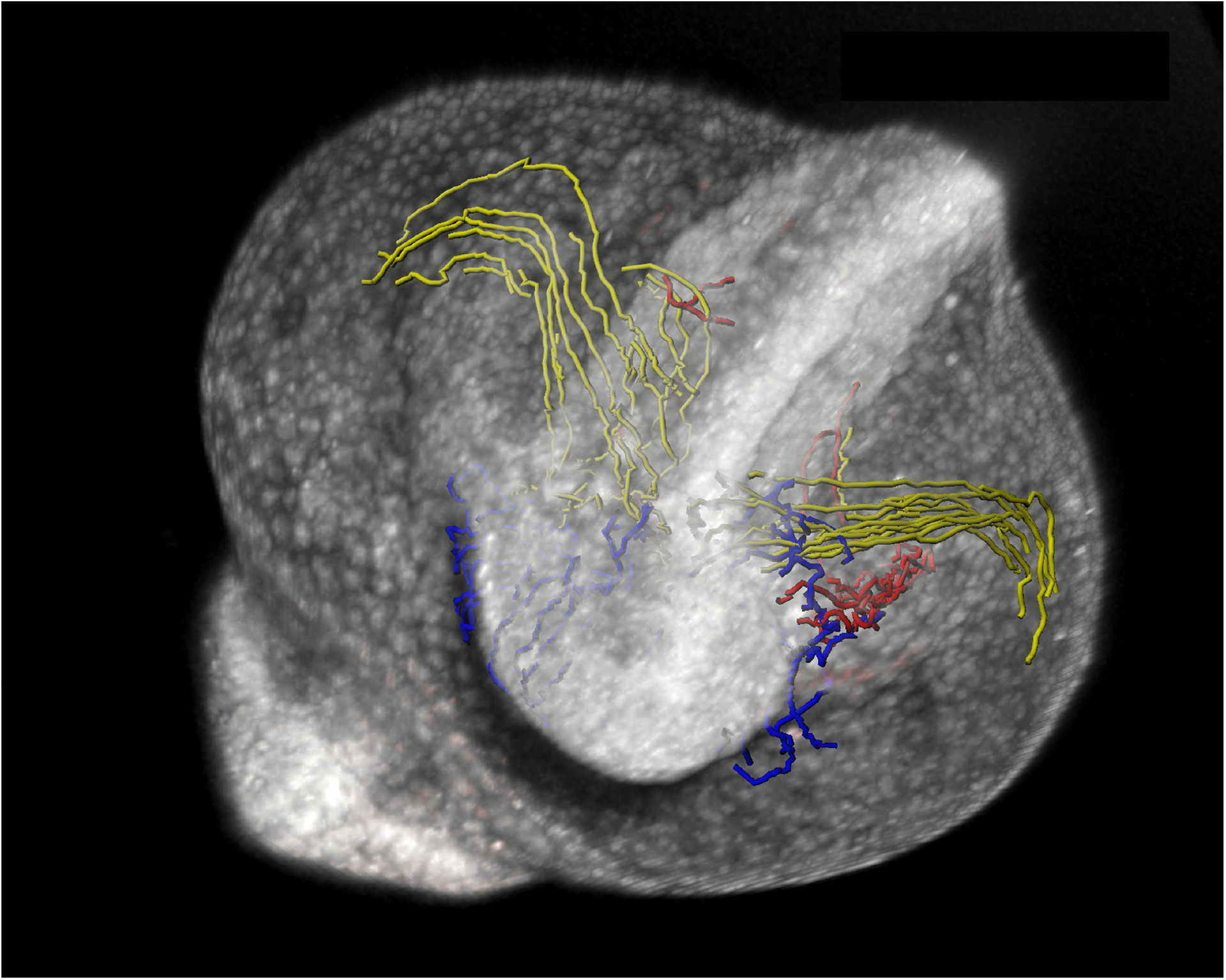
Relative proportional volumes of the spinal cord, paraxial mesoderm and notochord. A-B) Confocal stacks of embryos at the 15SS were used to build surface reconstuctions of the spinal cord (blue), paraxial mesoderm (green) and notochord (purple). C) Proportional volume of each tissue with respect to whole posterior body volume. Measurements are from three independent embryos. **Cover suggestion. This image shows the results of live cell tracking of posterior body contributions from 90% epiboly to the 12 somite stage.**

### Methods

#### Preparation of zebrafish samples

Wild-type AB embryos were obtained from zebrafish (Danio rerio) lines maintained following standard procedures. Embryos were injected at the one-cell stage with either 500pg of Kikume RNA (Hatta et al., 2006), a nuclear targeted version of the construct (nls-Kikume; kind gift of Ian Scott) or with a 500pg nls-mCherry together with 500pg membrane-GFP. Embryos were raised at 28°C and staged according to number of somites. For live imaging, 12 somite stage embryos were embedded in a small drop of 1% low melting point agarose within the central glass ring of a glass-bottomed petri dish (Mattek; 35mm petri dish, 10mm micro well. No. 1.5 cover glass). The agarose surrounding the tailbud was then removed using a pulled capillary tube.

#### Cell tracking and mapping of tailbud boundary

Embryos previously injected with nls-Kikume were mounted at 90% epiboly in 1% low melting point agarose and imaged on an inverted Leica SP5 confocal microscope with a 20X air objective. Small regions of around 10 to 20 nuclei were photo-converted from green to red by exposing the cells to 10 200Hz scans of a 405 laser at 60X zoom. Red nuclei were tracked in 4D using the image analysis tool Imaris (Andor technologies). The resulting tracks were filtered to include only long tracks lasting from 90% epiboly through to the 12 somite stage and manual corrected for tracking errors. The tracks were then colour coded depending on their final position at the 12-somite stage. Surface renderings of the entire embryo were automatically segmented on the green channel using the ‘surface’ tool in Imaris, and reference points were added at 50μm intervals along the embryonic midline from the most dorsal part of the blastopore at the 3 somite and 6 somite stages. Two lines were measured from these reference points to tracks at a boundary between either the tail-fated cells and the non-axial epidermis, or between tail-fated cells and trunk-fated cells. 3-and 6-somite staged reference embryos previously imaged by light-sheet microscopy were used to map the full complement of boundary points with the use of midline reference points as described. Surface reconstructions of the prospective tail region from one half of the embryo were then created by manually drawing contours at 10μm intervals.

#### Volume segmentation and morphometric analysis

Segmentation of tail volumes was performed using the manually create surface function of Imaris. Contours were manually drawn around the edge of the posterior body at 10μm intervals in z, bordering on the last formed trunk somite boundary of each species (somite 12 for zebrafish, 33 for mouse, 40 for lamprey and 20 for dogfish (Richardson et al., 1998). Surface reconstructions were then automatically constructed using these contours, allowing for the extraction of volume measurements. Measurement points were placed at every other somite boundary along the axis, and the values summed to give the length of the axis at any given stage. Height and length measurements were made at each length measurement point and plotted as described in the main text.

For photolabelled experiments, surface renderings were automatically created based on the red channel at consecutive stages through the movies. Volume, height and width measurements were made as described above for whole tail segmentations.

#### Preparation of mouse, dogfish and lamprey samples

Mice: mT/mG double-fluorescent Cre reporter mice were utilized in t order to generate mice with membrane targeted mTomato to visualize membranes (Muzumdar et al., 2007). Dogfish (*S. canicula*) eggs and lamprey embryos were produced by the Roscoff Marine Station and incubated at 17°C in oxygenated sea water until dissection and fixation in 4% PFA. Embryos were staged according to (Ballard et al., 1993). Lamprey (*L. fluviatilis*) embryos were obtained by in vitro fertilization and incubated at 17°C in oxygenated tap water until the required stages. Staging was performed according to (Tahara, 1988).

#### Imaging of fixed samples

Fixed zebrafish embryos previously injected with nls-mCherry and membrane-GFP were embedded in 1% low melting point agarose an imaged either on a Zeiss Lightsheet Z.1 microscope (at 3-and 6-somite stages) or on a Leica SP5 one-photon confocal at subsequent stages. Lamprey, mouse and Dogfish embryos were incubated with DAPI in PBS-0.01% Tween overnight at 4 °C prior to imaging. Lamprey embryos were using a Leica SP5 one-photon confocal. Mouse and Dogfish embryos were mounted in small chambers containing RapidClear solution (Sunjin labs) and imaged on a Zeiss LSM 7 MP multiphoton microscope. Images were tiled and automatically stitched together using microscope software in order to visualize the entire posterior body of large specimens.

## Acknowledgments

We would like to thank Alfonso Martinez-Arias for critical readings of the manuscript and Eric Theveneau, Simon Restrepo and Carlos Carmona-Fontaine for further essential commentary. We are indebted to Pascal Dardenne for excellent animal care and Christine Chevalier for help in isolating mouse embryos. We would further like to thank Ian Scott for the gift of the nls-Kikume construct. Finally, we acknowledge the Plateforme d’Imagerie Dynamique, Institut Pasteur, for their excellent imaging service and support. The authors declare no competing interests.

## Author Contribution

BS, EH and JFN conceived the project. BS performed most of the experiments. FD aided in performing morphometric measurements. EH performed live imaging for gastrula stage fate maps. RL and SM aided in staging and obtaining lamprey and dogfish embryos. BS and EH interpreted the results. BS wrote and EH edited the paper prior to submission.

